# Hi-C assembled genomes of estuarine populations reveal virus-microbe associations and a broad interaction range of a cyanophage

**DOI:** 10.1101/2023.12.06.570405

**Authors:** Christina Rathwell, Cedar McKay, Gabrielle Rocap

## Abstract

Aquatic microbes play key roles in global biogeochemical cycles and their viral-induced mortality influences the flow of carbon and nutrients between the dissolved and particulate pools. However, many microbes remain uncultivated, hindering understanding of their metabolic capabilities and preventing isolation of viruses that infect them. Here we augment metagenomic sequencing with Hi-C, a proximity-linkage method whereby DNA within a cell is physically bound and then sequenced to link contigs within a metagenome that originated from the same cell. In a size-fractioned water sample from beneath the euphotic zone in a hypoxic estuarine fjord in Puget Sound, WA we resolved 49 proximity-linked bins above 50% complete, including 21 Hi-C Assembled Genomes (HAGs) over 90% complete and a nearly complete genome of the eukaryotic green alga *Picochlorum*. Viral and microbial sequence within the same HAG identified 18 virus-microbe interactions. A myovirus and a siphovirus were associated with 2 different genera within the Saltatorellus clade of Planctomycetes, a phylum for which no virus has been identified. A partial *Phycodnaviridae* genome linked to Haptophyte sequence is consistent with contemporaneous observations of a dissipating coccolithophore bloom. A cyanophage S-CAM7-like sequence had a broad interaction range. It was associated with a partial *Synechococcus* genome in the >3.0 µm size fraction and with a Gammaproteobacteria related to *Alcanivorax* in the 0.2µm-3.0µm fraction. We suggest that viruses produced in surface waters that are shuttled to depth on sinking aggregates may interact with different hosts in deeper waters, providing an important avenue for gene transfer across broad taxonomic ranges.

**Importance:** Aquatic microbes are important in global elemental cycling. Knowing which viruses infect them in the environment remains a challenge. Using Hi-C, a molecular technique to physically link DNA within a cell, we assembled nearly complete genomes of both prokaryotes and eukaryotes from a hypoxic estuary. Hi-C links captured virus-host interactions for known virus-host pairs and for hosts with no previously known viruses. The same virus was linked to two distinct microbes in different size fractions of water, suggesting it has a broad host range. Viral lysis in surface waters generates sinking particles that deliver newly produced viruses to deeper waters where they interact with different potential hosts, providing an opportunity for gene exchange between unrelated microbes.

## Introduction

Marine microbes influence global biogeochemical cycles through carbon fixation, oxygen production and the sequestration of carbon to the deep ocean via the biological pump^1,2^. Within microbial communities, viruses play a large role in the distribution and composition of organic material^3^. Both heterotrophic and autotrophic microbial populations experience infection and undergo lysis, redistributing nitrogen- and phosphate-depleted organic material into dissolved pools in the viral shunt^4–6^. However, viral infection and cell lysis can also cause increases in particle aggregation and sinking, strengthening the biological pump in the viral shuttle^7–9^. Viruses alter the genomic potential of microbial communities as agents of density-dependent mortality of specific hosts^6,10–12^ and as vehicles for the exchange of genetic material^13,14^. Furthermore, a virocell expresses both viral and microbial genes, resulting in altered functions and contributions to biogeochemical cycling^15–17^.

Unfortunately, the majority of environmental viruses remain uncultivated and uncharacterized. The fraction of viruses that have been characterized through cultured isolates can be identified in the environment for presence, adsorption, infection rates, and transcription, which can lead to insights into viral ecology^18–21^. In the absence of an isolated reference, growing amounts of metaviromic sequence data^22^ have demonstrated the vast genetic diversity of viruses but are less well able to connect them to specific biogeochemical functioning. A critical remaining need is to determine which viruses are interacting with which hosts in the environment, in order to fully define the lifestyles and behaviors of viruses within different environmental contexts.

Isolation of novel viruses relies on having a cultivated host, yet approximately 81% of all microbial cells on Earth are from an uncultivated class or genus^23^. In the marine environment, although some of the most abundant organisms are in culture, single amplified genomes (SAGs) and metagenome assembled genomes (MAGs), make up over 40% of the genomes available for microbial species in the Genome Taxonomy Database, and these techniques are now commonly used to characterize uncultivated communities^24–35^. SAGs sample a very small volume, can identify microscale variation within a bacterial population, and can elucidate virus-host interactions for individual cells^36^. MAGs are derived from bulk environmental samples and provide population level characteristics of communities^37^. However, MAGs are unable to link viral genomes with host genomes as sequences are binned through nucleotide signatures such as tetranucleotide frequency and sequence coverage across samples, both of which may differ more between viruses and their hosts than between microbial populations^37^.

High throughput chromosome conformation capture (Hi-C)^38^ is another promising technique for binning sequences within a metagenomic assembly and can also link mobile genetic elements, such as viruses, with host metagenomes^39–41^. The Hi-C method involves crosslinking DNA in a live population using formaldehyde to hold DNA shape in place. Next, DNA is sheared and ligated in a dilute reaction, allowing DNA that was proximal in a cell but perhaps distant along the linear genome sequence, or even from a separate DNA molecule, to be joined together. These junctions are labeled, concentrated, and sequenced. This approach results in short DNA sequence reads that are composed of two different regions of DNA which were physically close to each other in the live cell. Hi-C has been used extensively to determine the scaffolding of chromosomes^42,43^, demonstrating how inter-chromosomal contacts might affect cell function^44^. This method has also been applied to cluster contiguous sequences (contigs) of a mixed culture metagenome into individual genomes^39–41^ and Hi-C has been extended to more complex communities to cluster over 900 MAGs from a rumen microbiome metagenome^45^ and 428 MAGs from sheep gut microbiomes^46^. Hi-C has also been used to investigate virus-host linkages within a cattle rumen microbiome^47^, to capture actively replicating viruses in the human gut microbiome^48^ and has detected broader host ranges of viral elements in a sheep gut microbiome than previously seen through cultured isolates^46^. Here we explore the potential of Hi-C in the aquatic environment, home to complex microbial communities that are much more dilute than mammalian gut microbiomes. We applied Hi-C linkages to two size-fractionated metagenomes in Hood Canal, Washington, and used a conservative Hi-C clustering approach to improve binning of microbial metagenomic sequences and identify specific viruses and the potential hosts they are interacting with in the environment.

## Results

### Sampling and Proximity Linkage-Enabled Bin Generation

Hood Canal is a long narrow estuary connected by a shallow sill to Puget Sound, Washington at its northern entrance (Figure 1A), resulting in relatively slow flushing times and seasonal hypoxia in the southern end. The water in southern Hood Canal (bottom depth 35 m) on August 15, 2016, was sharply stratified with a shallow (<10 m) mixed layer and chlorophyll maximum at 5 m; water at the sampling depth of 25 meters was hypoxic (0.5 mg/L, or 15.5 µmol/kg^3^; Figure 1B-C). Continuous data collected by the Twanoh Bay ORCA mooring in Hood Canal indicated that the bottom waters had been hypoxic for over a month (Figure 1B). Additionally, earlier in July a coccolithophore bloom, visible from space, had established in Hood Canal (Figures 1A and S1), and was visually confirmed to be present at time of sampling. Virus like particle concentrations at 25 m were 8.76 x 10^8^ ml^−1^, which was 18.4 times that of microbial cells (4.77 x 10^7^ ml^−1^) (Figure 1D). Independently size-fractionated samples from 25 m were used to generate both Hi-C sequence libraries and standard short read metagenomic sequence libraries (Table S1).

**Figure 1.**
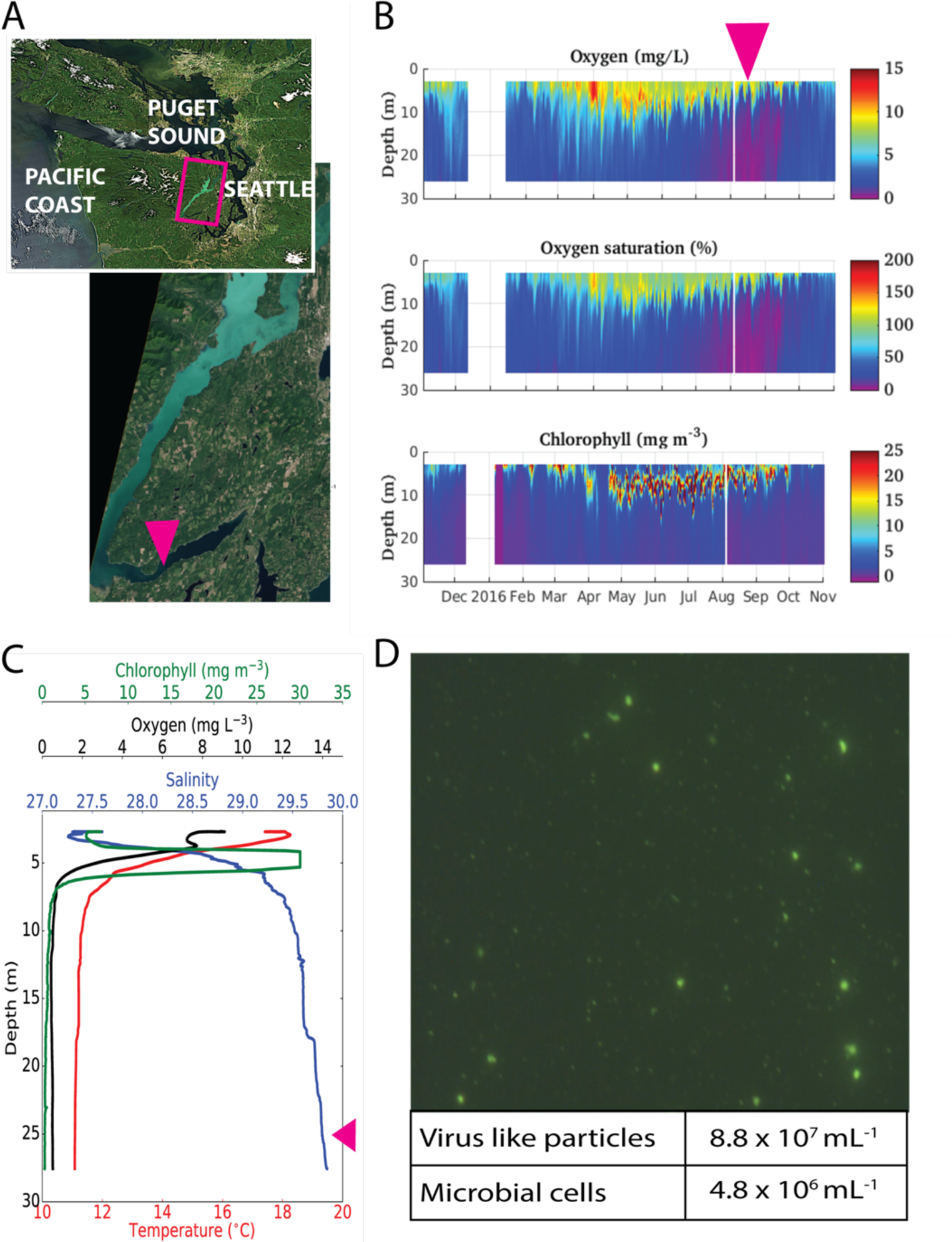
Environmental context. **A**) Satellite imagery from July 26^th^, 2016, downloaded from NASA MODIS archive, showing turquoise coccolithophore bloom spreading from the north of Hood Canal into the southern reaches of Hood Canal. Pink rectangle and inset map indicates location of Hood Canal. Pink arrow indicates mooring location and sampling site. **B**) Annual data from ORCA mooring located in Twanoh Bay, Hood Canal showing twice daily oxygen concentrations, oxygen saturation and chlorophyll concentrations with depth for 2016. Pink arrow indicates time of sampling. **C**) Depth profile of salinity, chlorophyll, oxygen and temperature from ORCA mooring on day of sampling, August 15, 2016. Pink arrow indicates depth sampled. **D)** Epifluorescent microscopy image of whole seawater at 25 m, with the average virus like particle and microbial concentrations at 25 m.

We used two different established methods to cluster assembled contigs based on proximity linkages as well as a linkage-independent method. MetaBat attempted to cluster all contigs greater than 1500 bp and generated 165 and 74 bins from 107,590 and 54,957 contigs in the 0.2–3 µm and >3 µm size fractions respectively (Figure 2A-B, Table S2). Proximity linkage-enabled binning employed only the subset of contigs that were part of a Hi-C read pair (37.5% of contigs in the 0.2–3 µm library, and 18.9% in the >3 µm library; Figure 2A-B Table S2). HiCBin generated 65 and 38 bins from 56,179 and 38,116 contigs in the 0.2–3 µm and >3 µm size fractions respectively. In contrast, the ProxiMeta algorithm which also utilized the Hi-C read pairs incorporated more contigs than HiCBin and resulted in more bins; 301 and 261 bins from 85,156 and 50,925 contigs in the 0.2–3 µm and >3 µm size fractions respectively. None of the ProxiMeta bins were more than 10% contaminated as determined by checkM, while HiCBin resulted in 17 bins that were greater than 10% contaminated and MetaBat generated 51 bins greater than 10% contaminated (Figure 2E-F).

**Figure 2.**
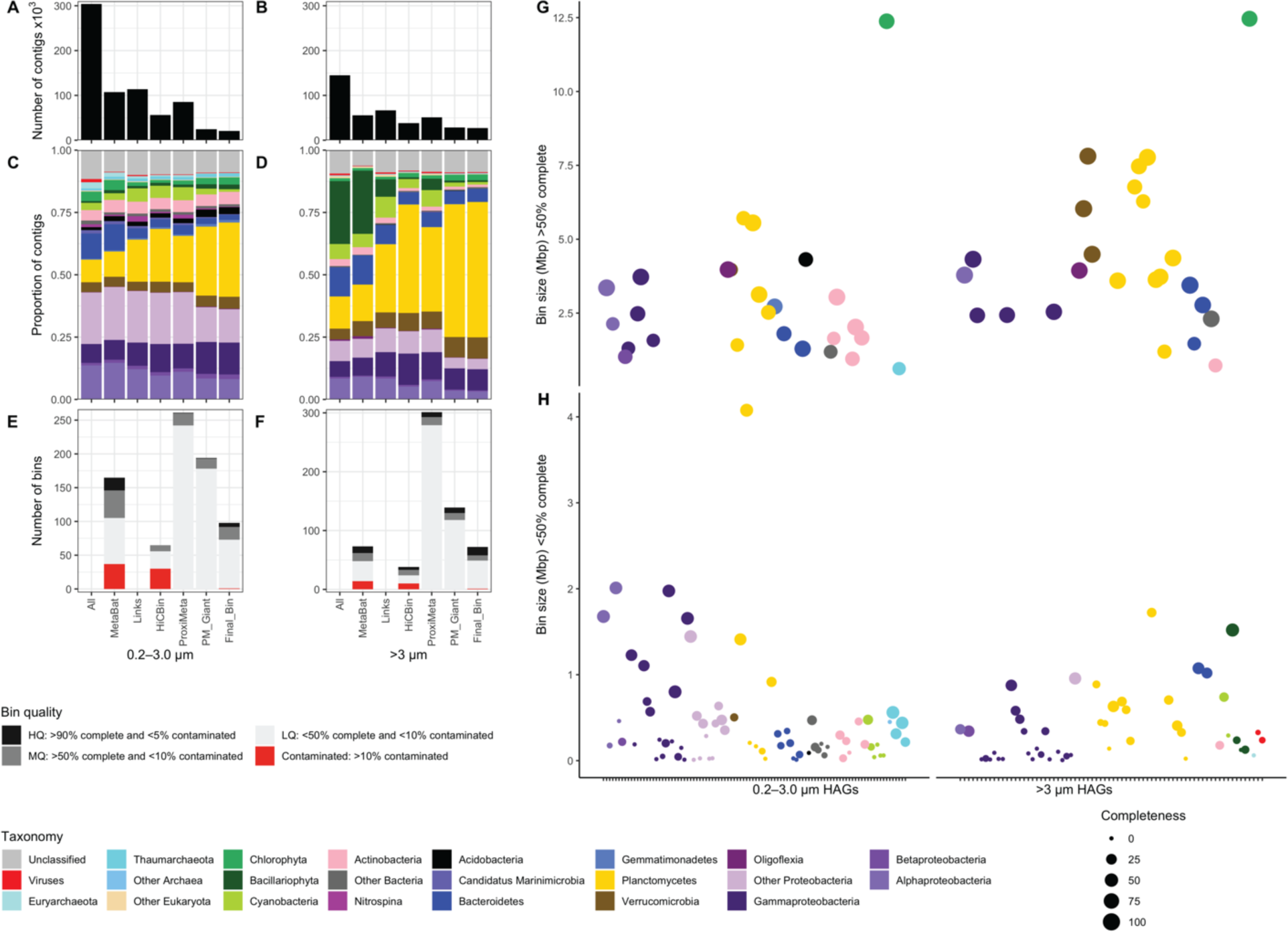
Overview of Hi-C linked bin generation and assessment. **A-B)** Number of contigs greater than 1 kbp for each step including: the initial assembly (All), bins generated without using Hi-C links (MetaBat), contigs containing a link to a Hi-C read (Links), bins generated by HiCBin (HiCBin), bins generated by ProxiMeta (ProxiMeta), ProxiMeta refined giant component clusters (PM_Giant), and the ultimate intersection of HiCBin bins with ProxiMeta refined giant component clusters (FinalBin). Values are shown for the 0.2–3.0 µm library (A) and the >3 µm library (B). **C-D)** Proportion of the contig pool included in each step above assigned to different taxonomy for the 0.2–3.0 µm library (C) and the >3 µm library (D). **E-F)** Summary of checkM data for each binning methods above for the 0.2–3.0 µm library (E) and the >3 µm library (F). Contaminated bins (>10%) are shown in red regardless of completeness and non-contaminated bins are shown in grey and black based on quality classification. **G-H)** Completeness, size and taxonomy of final proximity linkage bins separated by those >50% complete (G) and < 50% complete (H). Each bin is represented by a dot color-coded by DIAMOND blast taxonomy. Dot radius is relative to checkM completeness. Note different scales for y-axis (bin size).

Because a central goal of this study was to connect viral sequences with their potential hosts, we used a conservative approach to generate the final bins, prioritizing stringency of proximity linkages over maximizing bin size. First, the ProxiMeta bins were further refined using a Giant Component analysis to retain only contigs that were part of the most connected web in each bin (Figure S2). Like the original bins, these ProxiMeta Giants were not contaminated above 10% (Figure 2 E-F). Because HiCBins were generally larger than ProxiMeta bins but more likely to be contaminated (Figure 2 E-F), we combined the more aggressive clustering of HiCBin with the strict filtration of the ProxiMeta clustering to create intersection bins (Figure S3) and thus enhanced completeness without having contaminated bins. This approach resulted in 170 final bins (98 in 0.2–3 µm and 72 in >3 µm), none of which were contaminated above 10%. Of those 170 bins, 21 are high quality draft genomes (>90% complete, <5% contaminated), 28 are medium quality (>50% complete, <10% contaminated) and the remainder can be considered low quality draft genomes (<50% complete, <10% contaminated; Figure 2 E-F, Table S2).

The overall representation of each taxonomic class among assembled contigs, linked contigs and contigs binned using the various methods was generally similar, although there were some notable exceptions (Figure 2 C-D). In both size fractions Planctomycetes represent a much larger fraction of contigs with Hi-C links than of total contigs: in the >3 µm library Planctomycetes represent 10% of total contigs assembled, 21% of contigs with links, and 41% of contigs in final bins (Figure 2D). In contrast, in the >3 µm library, Bacillariophyta contigs made up 25% of assembled contigs, but only 7% of linked contigs and only 1-5% of any subsequent linkage-dependent binned contigs (Figure 2D). The 170 final intersection bins were assigned to 19 different phyla with the majority of bins identified as Proteobacteria (70 bins) or Planctomycetes (34 bins) (Figure 2 G-H). Bins greater than 50% complete spanned 13 different phyla assignments.

### High Quality Hi-C Assembled Genomes

The 21 Hi-C assembled genomes (HAGs) that were >90% complete and <5% contaminated, included 7 Planctomycetes, 6 Proteobacteria, 3 Verrucomicrobia, 3 Actinobacteria, and 2 Bacteroidetes (Table S3). Notably, the 6 bins classified as Proteobacteria represented 3 pairs from different orders (*Oligoflexales*, *Cellvibrionales* and *Rhodobacterales*) in which a highly similar genome (>99% ANI within each pair) was independently assembled in each size fraction (Figure S4). The *Oligoflexales* (IB_0.2_006 and IB_3.0_013), whose class *Oligoflexia* has recently been reclassified as a new phylum *Bdellovibrionota*^49^, each had a best blast hit to the predatory bacterium *Pseudobacteriovorax antillogorciicola*. The *Cellvibrionales* bins (IB_0.2_005 and IB_3.0_006,) were members of the genus *Halioglobus* and the *Rhodobacterales* bins (IB_0.2_003 and IB_3.0_004) were classified as *Rhodobacteraceae*.

High representation of Planctomycetes in contigs with Hi-C links enabled assembly of 7 high quality draft genomes and 7 medium quality draft genomes that represent at least 8 distinct Planctomycetes populations from five major clades including *Pirellula*, *Bythopirellula*, *Gimesia*, Phycisphaerae, and the newly cultured *Saltatorellus* clade^50^ (Figure 3). Within the Planctomycetes there were also cases of independent assembly of closely related genomes in each size fraction, including one pair most closely related to *Mariniblastus fucicola* (IB_0.2_001, IB_3.0_005, 99.4% ANI), as well as a pair in the *Bythopirellula* clade (IB_0.2_007, IB_3.0_010, 99.2% ANI; Figure 3, Table S3). Proximity linkages also enabled lineage resolution of closely related taxa in the same size fraction in the *Saltatorellus* clade (IB_3.0_008, IB_3.0_014). Phylogenetic placement of concatenated core Planctomycetes genes from the HAGs frequently places them closest to MAGs from metagenomic sequencing in the Mediterranean Sea^24^ (named TMEDX in Figure 3). This placement is not surprising, as many cultured Planctomycetes isolates come from populations present in freshwater, soil, peat moss and even plumbed water, and may not be the best representatives of marine populations^51^. There were no bins of any quality identified as members of the *Candidatus* Brocadiaceae clade, which contains organisms capable of performing anammox.

**Figure 3.**
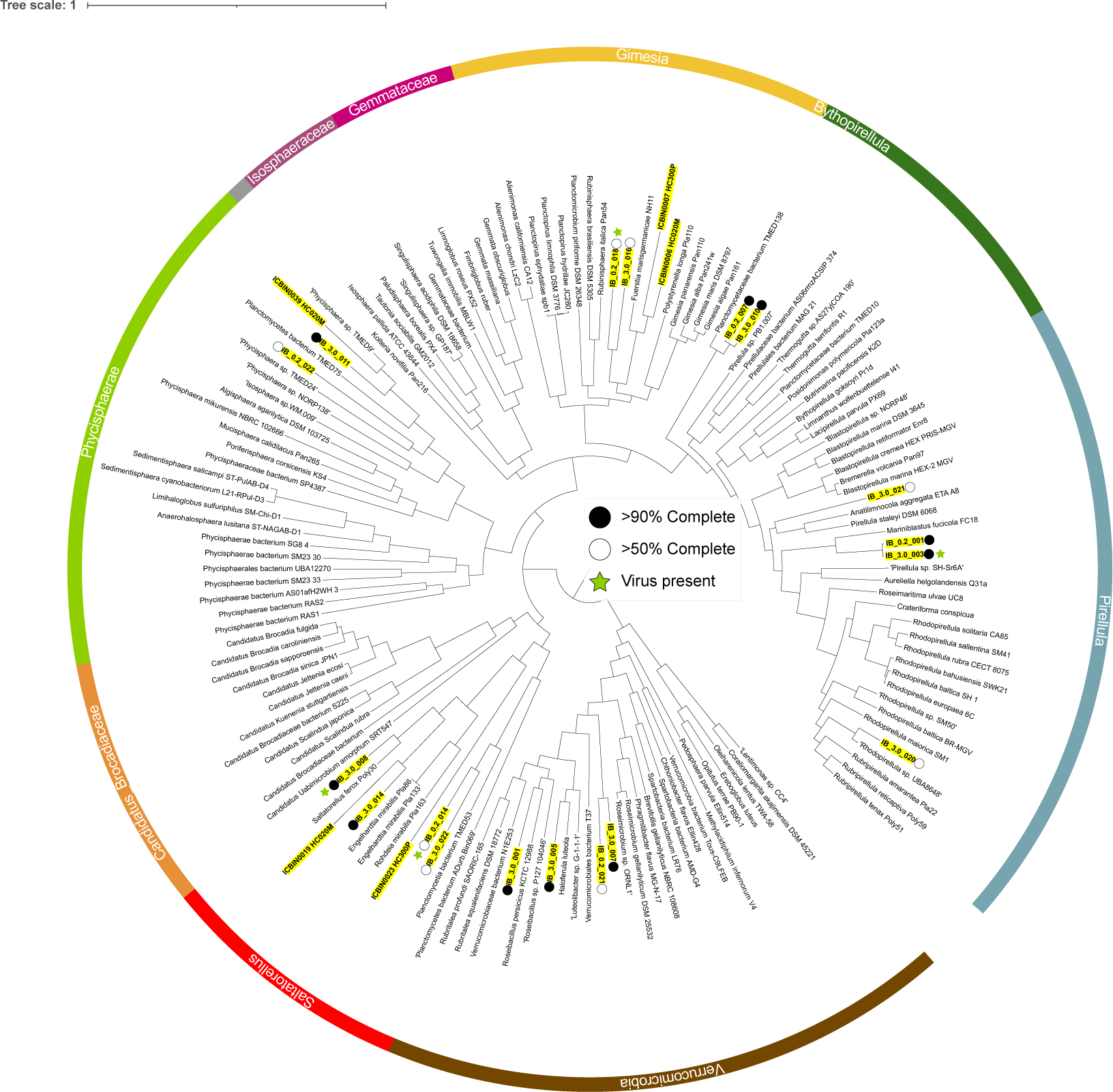
Maximum likelihood phylogenetic tree of Planctomycetes and Verrucomicrobia based on concatenated proteins of 571 single copy genes from the Planctomycetes lineage. The 18 Hi-C assembled genomes >50% complete are indicated as medium quality (MQ; 50-90% complete, 8 HAGs) or high quality (HQ; >90% complete, 10 HAGs). Viral sequence present within a HAG is indicated by a green star. The Verrucomicrobia were used as the outgroup. Numbers at nodes represent % of Felsenstein bootstrap values (100 bootstrap replicates were performed, at which point the replicates were assessed to have converged).

In addition to the 21 high quality draft genomes of prokaryotes, two ∼12.4 Mb bins, one from each size fraction (IB_3.0_017, IB_0.2_010), each contained a nearly complete genome of *Picochlorum,* a unicellular green alga from the phylum Chlorophyta (Figure 2G). Phylogenomic analysis indicates that they are closely related to each other (99.86 % ANI) and to *Picochlorum* NBRC102739 (Figure S5). Although these two bins were judged as 47–77% contaminated using checkM, which employs a marker set intended for bacterial and archaeal genomes, analysis using BUSCO^52^ with a set of 1519 Chlorophyta specific gene markers suggests that they were approximately 86% complete with less than 1% of the single-copy orthologs duplicated. This completion statistic is in the range of complete published *Picochlorum* genomes from cultured isolates, 68–96% (Table S4) and may be an underestimate, as the gene prediction on these contigs using prodigal likely failed to fully capture accurate eukaryotic gene models. Thus, as with the three pairs of Proteobacteria and two pairs of Planctomycetes, proximity linkages were able to independently assemble nearly complete and nearly identical eukaryotic genomes from the two size fractions.

### Proximity Linkages Identify Viral Host Associations

Virus-host associations were examined by searching for viral sequences within the final proximity linkage bins, which were generated with a conservative binning approach to minimize contamination (Figure S3). We identified 18 bins that contained >10 kb of viral sequence and that could be classified with DemoVir (Table S5, Figure 4). Viruses identified included *Myoviridae* (7)*, Siphoviridae* (3), *Podoviridae* (2), unclassified Caudovirales (2) and *Phycodnaviridae* (4). CheckV determined two of these viral sequences to be potential prophage (Figure 4). We used the taxonomy of the non-viral sequence in each bin to determine the microbial populations with which each viral genome interacted. The HAGs containing viral sequence varied in size and completeness of both host and viral genomes and included five of the high-quality draft genome bins as well as six bins that had minimal host genomes (3% complete or less; Figure 4, Table S3). The majority of the viral contigs within HAGs were not placed in bins by the Hi-C independent binning method MetaBat2 (Figure S6)

**Figure 4.**
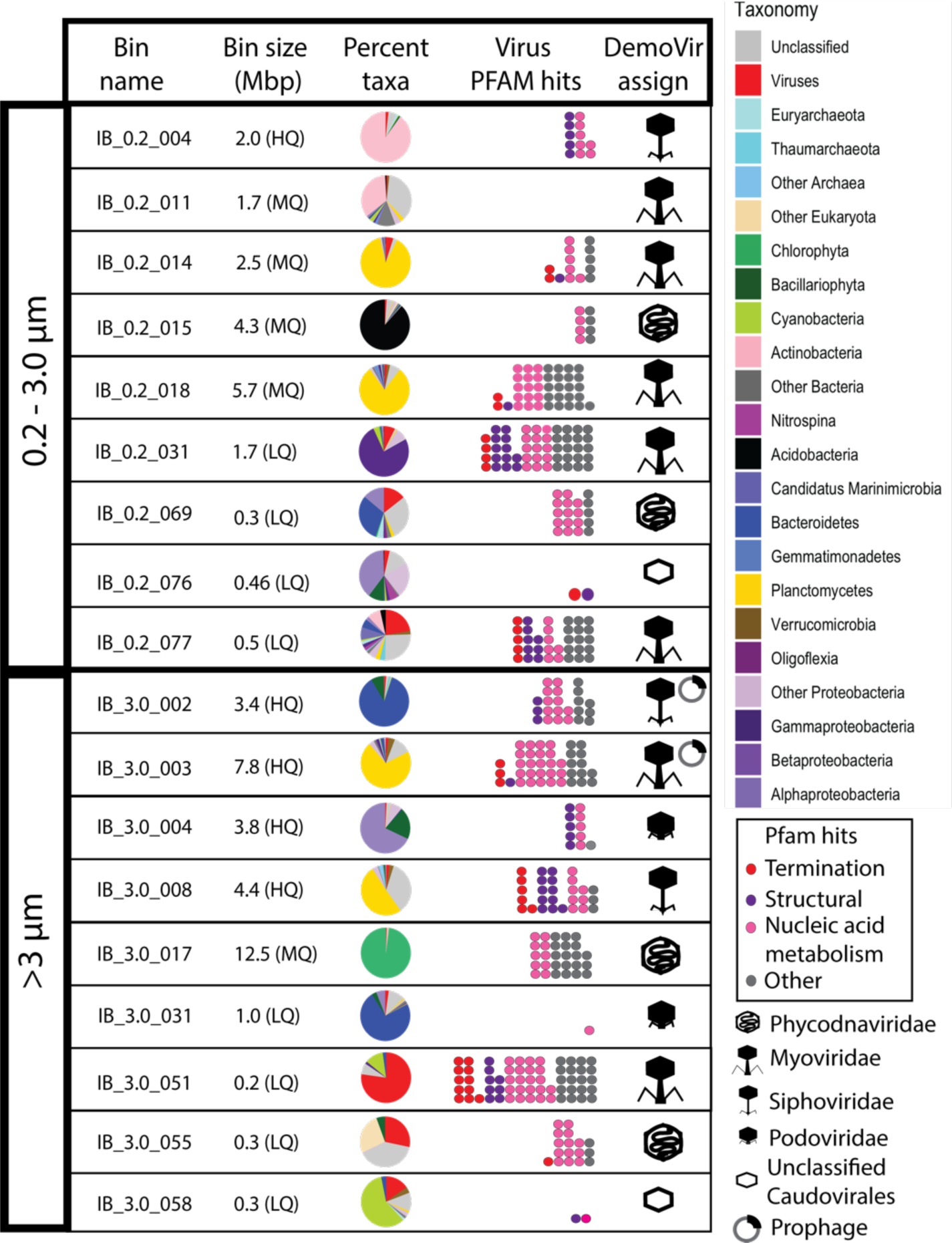
Virus-microbe interactions detected in HAGs. Each row contains data about the bin in which the virus sequence was clustered in and the virus sequence itself. Bin information includes name, size, and quality (where high, medium and low quality determination is indicated by HQ, MQ and LQ, respectively) and pie chart representing percentage of total sequence length assigned taxonomy by DIAMOND blast. Virus information includes dot plot of color-coded circles indicating viral proteins identified by Pfam search and icon with DemoVir viral taxonomy assignment.

Although viruses infecting marine or estuarine Planctomycetes have not been previously identified, 4 Planctomycetes bins ranging in size from 2.5 Mb to 7.8 Mb contained viral genomes (Figures 3 and 4). In the 0.2–3 µm size fraction two *Myoviridae* viruses with many viral genes were found in IB_0.2_018 (*Gimesia* clade) and IB_0.2_014 (*Saltatorellus* clade) (Figures 3 and 4). IB_0.2_018 contained genes encoding RNA polymerase, portal protein, several peptidases, nuclease, a major capsid protein, lysins and DNA polymerase (Figure S7). IB_0.2_014 had fewer canonical viral genes but included genes encoding the terminase large subunit, a major capsid protein, DNA recombination, helicases and an integrase (Figure S7). In the >3 µm size fraction IB_3.0_008 is a near-complete HAG related to *Saltatorellus ferox* containing *Siphoviridae* including genes for many structural and assembly proteins, such as major capsid protein, portal protein, 6 tail proteins, tape measure protein, terminase and head assembly protein (Figure S7, Table S6). Finally, also in the >3 µm size fraction, a potential prophage was identified in IB_3.0_003, a near-complete HAG closely related to *Mariniblastus fucicola*. This HAG contained a *Myoviridae* sequence including genes for Thy1, multiple terminase units, tail protein, RNA polymerase, portal protein, DNA polA, and more (Figures 4 and S7). Although Pfam searches conclusively identified canonical viral genes in all four of these bins, Blastn searches against viral RefSeq resulted in very low identity matches, generally to non-marine viruses (i.e., *Bacillus phage, Pseudomonas phage*, *Caulobacter phage*), suggesting that these are novel viruses with no reference genomes sequenced yet.

Four of the 18 identified viruses, all from the >3 µm size fraction, were associated with photoautotrophic hosts, even though the sampling depth of 25 m was below the euphotic zone (Figure 1C, Figure 4). A *Phycodnaviridae* was identified in the *Picochlorum* bin IB_3.0_017, and a Caudovirales was identified in a very incomplete *Synechococcus* bin (IB_3.0_058) that contained nearly 10% viral sequence (Table S3, Figure 4). A third bin, IB_3.0_05, contained majority viral sequence, a *Phycodnaviridae* with 3 viral contigs totaling 92.8 kb. A smaller fraction of this bin sequence was identifiable as microbial in origin and included a plurality of Haptophyte contigs (6 contigs, 88.3 kb) classified in 3 different orders (*Isochrysidales, Prymnesiales* and unclassified Haptophyta) as well as 4 unclassified contigs (81.2 kb) and 4 others that were identified as some other Eukaryote, each a different one (51.4 kb). Genes on viral contigs in IB_3.0_055 most frequently had a blastx against nr best match to *Emiliana huxleyii* virus *EhV86*. However, percent identities ranged from 23% to 47%. Overall, these results suggest the presence of a haptophyte and its associated virus, both of which are only distantly related to sequences currently in the database. Their presence is consistent with contemporaneous observations of a coccolithophore bloom occurring in the sampled waters (Figures 1A and S1).

The fourth photoautotroph and virus bin, IB_3.0_51, was also predominantly viral and contained a *Myoviridae* (98.9 kb, 69 contigs) most closely related to the myovirus, *Synechococcus* phage S-CAM7, which has a 214 kb genome and was isolated off the Southern California Pacific Coast on *Synechococcus* strain WH7803^53^ (Figure 5). Also in this bin are 21 contigs (80.1 kb) most closely related to clade I *Synechococcus* strain ROS8604 (Figure S8). As observed generally with viral sequence with HAGs (Figure S6), both HiCBin and ProxiMeta were able to pair *Synechococcus* contigs with S-CAM7-like viral contigs, but MetaBat2 was unable to place most of the S-CAM7-like viral contigs into any bin. Though this bin was very incomplete, there were 5 other incomplete clade I *Synechococcus* bins present across both size fractions, ranging in size from 15 kb to 160 kb, with a checkM completeness of 0–6% (Table S3, Figure S8). Furthermore, the viral sequence in IB_3.0_51 was one of two of the viruses identified that had more than 50 genes for canonical viral proteins as determined by Pfam searches (Figures 4 and S7), and possessed genes for tail proteins, portal proteins, phage lysozyme, peptidases, neck proteins, major capsid proteins, terminase, helicases, head assembly proteins and baseplate proteins.

**Figure 5.**
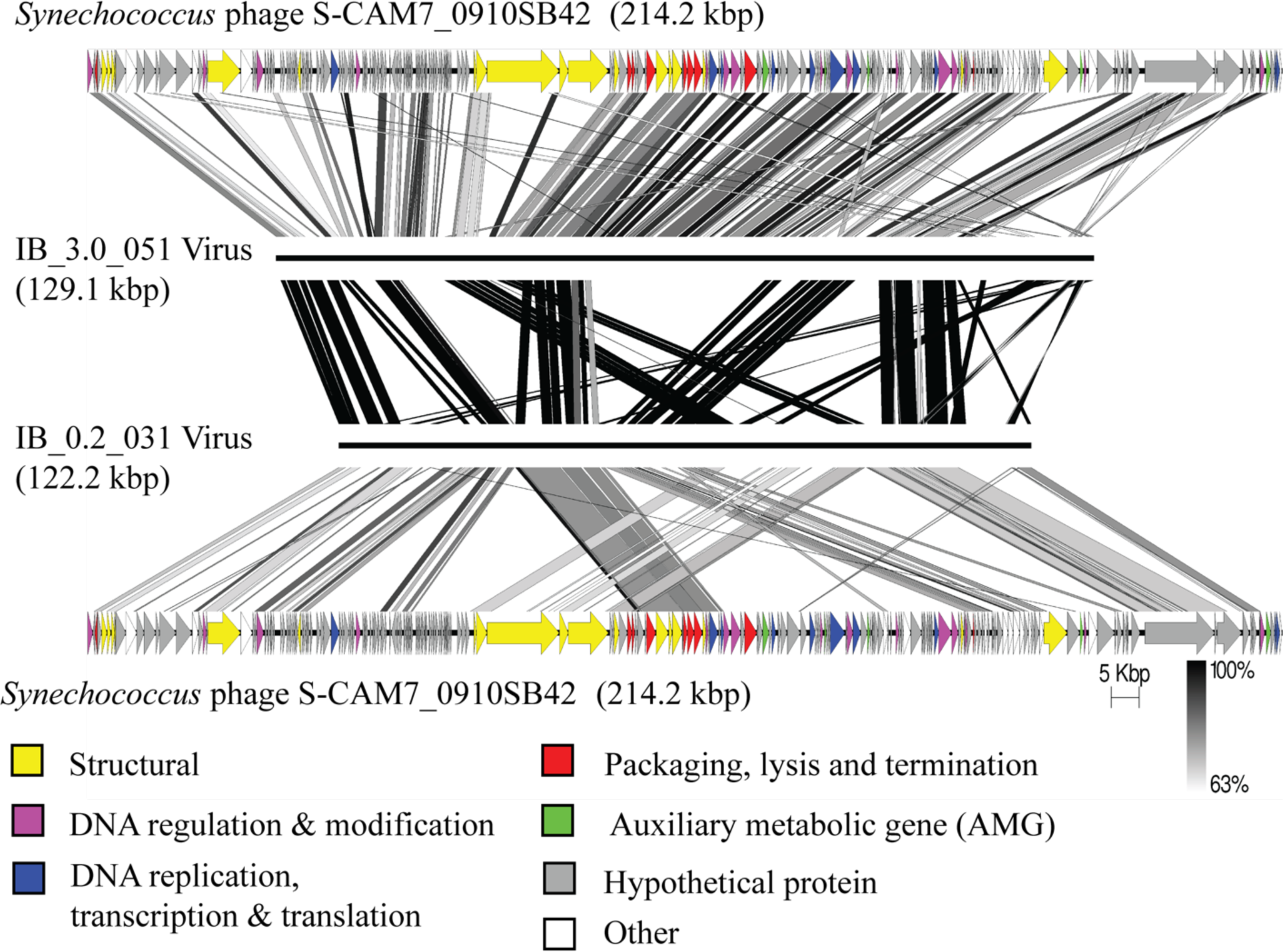
S-CAM7 virus detected in both size fractions interacting with different hosts. Genome similarity between the *Synechococcus* phage S-CAM7 0910SB42 published genome (top and bottom rows) and viral genomes from bins IB_3.0_051 (*Synechococcus*) and IB_0.2_031 (*Alcanivorax)*. Virus genes are annotated on the S-CAM7 genome and color coded by function. Scale bar represents percent nucleotide identity between each pair of genomes. See table S7 for list of AMGs in each genome.

An additional viral genome closely related to *Synechococcus* phage S-CAM7 was identified in the other size fraction (0.2–3 µm); however, this *Myoviridae* genome (122.2 kb, 9 contigs) was present in a bin (IB_0.2_031) containing 1.65 Mb of a Gammaproteobacterium most closely related to *Alcanivorax* (Figures 4 and 5). The viral sequence within this bin also contained more than 50 canonical viral proteins, most with high percent identity (63–100%) to both the S-CAM7 0910SB42 genome and to the S-CAM7-like viral sequence in IB_3.0_51 (Figures 5 and S7). Both S-CAM7-like viral genomes contain auxiliary metabolic genes (AMGs) previously described in cyanophage (Table S7). Notably, each virus contained the photosynthesis-related AMGs for *cp12*, a gene responsible for inhibiting the Calvin Cycle during infections in the light^54^, and *cpeT*, a bilin lyase, important for pigment adaptation in the freshwater cyanobacteria *Fremyella diplosiphon*^55^. These two genes are commonly reported in cyanophage genomes but are not present in the S-CAM7 0910SB42 genome^53^.

## Discussion

### HAGs and Virus-Host Interactions Made Possible Through Hi-C Sequencing

Despite their significant impact on biogeochemical cycles most members of aquatic microbial and viral communities remain uncultivated^22,23^. In order to characterize uncultivated microbial and viral populations and their intracellular interactions in the environment, we generated Hi-C proximity linkage libraries in tandem with bulk metagenome samples from two size fractions in the deeper, more saline layer of Hood Canal estuary in Puget Sound, WA. Using a stringent approach to linking contigs with Hi-C read pairs, we generated 49 bins greater than 50% complete and 21 bins qualifying as high-quality draft genomes^56^. Among the high quality HAGs were 5 pairs of closely related genomes (>99% ANI within each pair) independently assembled in each size fraction, providing an unintended replication of the method. In addition, two nearly complete genomes of the green alga *Picochlorum*, one in each size fraction, demonstrate the potential of Hi-C for assembling genomes of marine eukaryotes. Hi-C was particularly successful in assembling genomes from the phylum Planctomycetes, perhaps because these cells have cytoplasmic invaginations that order their genomes in condensed, membrane-bound, nucleoid structures^57^. The generation of over 20 high quality draft genomes from one sampling event shows the power of using proximity-links in environmental metagenomics. Enhanced assemblies will help elucidate the genomic potential of uncultivated microbes and can help bring more microbial taxa into culture, expanding available hosts for isolation of marine viruses^58^.

We specifically use the term HAG for the proximity linkage bins, to differentiate them from MAGs, since the Hi-C links enable binning of non-contiguous sequence (such as distinct chromosomes or plasmids) and even unrelated genomes that co-occur within the same cell. Indeed, some proximity linkage bins contained both viral and microbial sequences, potentially indicating the presence of viruses within cells, which ultimately allowed for designation of virus-microbe associations *in situ* for 18 HAGs. The majority of the viral contigs within HAGs were not placed in bins by Hi-C independent binning methods. Although Hi-C sequencing is a promising window into virus-microbe interactions in the environment, some caution is warranted interpreting sequence from HAGs. First, as with all methods aiming to detect viruses based on sequence, viral and microbial sequence can be hard to disentangle based on similarity to reference databases, because such a small fraction of viral populations has been isolated. Second, shorter contigs may not allow distinguishing between an infecting virus and a prophage integrated in the genome or present on a plasmid, as all will be linked with the microbial HAG using Hi-C reads, as observed in two cases here. Finally, as one HAG represents a community of microbial cells, if multiple viruses were infecting a given population at a time, one HAG could potentially contain multiple viral genomes. Closer inspection of viral sequences found within HAGs, including DemoVir assignment, Pfam assignment of genes, examination of single copy canonical genes and alignment of genomes, was necessary to identify the 18 virus-microbe associations here. These included four Planctomycetes HAGs ranging in size from 2.5 to 7.8 Mb containing viral genomes (3 *Myoviridae* and a *Siphoviridae*), the first potential identification of viruses infecting marine Planctomycetes, demonstrating this method’s ability to assemble viral genomes for greatly under characterized viral populations.

### Phytoplankton and their Viruses Delivered to Aphotic Depth on Particles

The viral shuttle hypothesis proposes that viral infection of surface primary producers contributes to particle flux to depth^59^. Coccolithophore blooms experiencing viral infection contribute heavily to particulate carbon flux to the deep ocean^8^, in part due to the ballast of their disc-like structures on the cell surface but potentially also due to the aggregation of cell material post-lysis^9^. Abundance of picophytoplankton such as *Synechococcus* and their viruses are also associated with generation of particles at depth^7^. In Hood Canal, at a sampling depth of 25 m, below the stratified surface layer and just 10 m above the seafloor, 4 bins in the >3 µm size fraction were composed of photoautotrophs associated with viruses, supporting the viral shuttle hypothesis^88^. Here the two bins containing both *Synechococcus* and cyanophage were in the >3 µm size fraction, instead of the 0.2–3 µm fraction where free-living *Synechococcus* might be expected. The assembly characteristics of some of these bins also appear to suggest a progressed lytic infection. The Haptophyta and one of the *Synechococcus* HAGs were predominantly viral, with very little host sequence present in the final bin, which might occur in the later stages of infection as viral genomes are replicated and host genomes are degraded^60^. To further support this hypothesis, model virus-host systems could be used in the laboratory to characterize the precise Hi-C linkage signal across an infection cycle with a view towards linking Hi-C sequencing data to stage of infection or type of viral interaction *in situ*.

### S-CAM7 Synechococcus Phage Interacts With a Broad Range of Hosts

The proximity links we obtained in this study identified previously characterized virus-microbe associations^53,61^ *in situ* while also revealing alternate virus-microbe interactions. In the particle-associated size fraction a cyanophage closely related to *Synechococcus* phage S-CAM7 paired with a clade I *Synechococcus*. S-CAM7 was isolated on clade V *Synechococcus* strain WH7803, but the ability to infect across clades of *Synechococcus* is not unexpected for a generalist cyanophage^62^. However, in the free-living fraction a virus even more closely related to S-CAM7 was linked to microbial sequence related to *Alcanivorax,* a genus of hydrocarbon-degrading Gammaproteobacteria^63^. Although this interaction may be surprising, there is some precedent for Proteobacteria to be interacting *in situ* with cyanophage, as SAG sequencing has identified cyanophages sequences related to S-CAM7 in *Roseobacter*^36^. Laboratory experiments have demonstrated that a generalist cyanophage may enter the cells of a resistant cyanobacterial strain and replicate the viral genome and assemble capsids, yet not complete a lytic infection^62^. The Hi-C linked sequences presented here cannot determine whether S-CAM7-like viruses in Puget Sound are producing a lytic infection in *Alcanivorax* populations but are evidence that the virus had entered these cells. Notably, S-CAM7 carries the fewest photosynthesis AMGs of cultured cyanomyoviruses^53^, which might occur if the evolutionary landscape of this virus included a broad range of interactions^64^. Future work assessing the ability of cultured S-CAM7 to enter non-cyanobacterial hosts and replicate or generate capsids is required to understand the evolutionary constraints on this viral population. Whether this S-CAM7 interaction with *Alcanivorax* represents a truly lytic infection or an aborted lytic infection, it provides an avenue for gene transfer via amongst diverse microbial populations.

To date the majority of aquatic microbial virus-host interactions have been witnessed *in vitro*, leading to the hypothesis that viruses in the environment have a narrow host range^65^. However, some laboratory evidence suggest viruses isolated on a single host population may increase their selectivity in future passages but that alternating hosts for the isolation increases viral host range, even if at an apparent cost to virus fitness^64^. The marine environment differs from cultures in many ways, particularly in its complex community composition, dilute environment and physical transport processes^22^. Viruses produced in surface populations may contribute to particle aggregation and be shuttled to depth where the primary host population is no longer abundant. Here viruses may interact with alternate species, allowing for genetic transfer across a broad range of taxa and potentially selecting for broad host ranges in viral populations^59,66^. Expansion of the use of Hi-C linkages and SAG sequencing^36,67,68^ to determine virus-host interactions *in situ*, and assessment of host ranges with cultured isolates will clarify the role of viral host selectivity in influencing both microbial population dynamics and carbon cycling and flux of carbon to depth in the ocean.

## Materials and Methods

### Collection Site and Environmental Conditions

Samples were collected from Twanoh Bay in Hood Canal of Puget Sound in Mason, Washington on August 15, 2016 from 4 pm to 6 pm. Sampling was done adjacent to an Ocean Remote Chemical Analyzer (ORCA) profiling mooring^69^, at 47° 22.5’N, 123° 0.5’W. The bottom depth at this site is at 35 m. Water was collected from 25 m depth with a Niskin bottle deployed from the deck of R/V *Mackinaw*, a vessel used to service the ORCA moorings.

### Epifluorescent Microscopy

Fifty milliliters of water were collected from 25 m and processed using established methods for enumeration via microscopy^70,71^. Water was treated with microscopy grade paraformaldehyde, immediately frozen on deck in liquid nitrogen, and stored back in the lab at –80°C until it could be processed further. The sample was thawed and filtered onto two 0.02 µm Anodisc filters. Dried filters were incubated in 2.5x concentration SYBR Gold for 25 minutes in the dark. Excess dye was removed, and filters were mounted on a slide using 50:50 Glycerol/1X TE. Samples were viewed at 1000x under excitation with a X-Cite 120 bulb. Images were captured of fields of view using the Nikon Eclipse 80i. Fluorescing regions were classified as microbial or virus like particles based on size and counted across 20 to 200 fields of view. Pixel to micrometer conversion was calculated using a 10 µm stage micrometer ruler at equal magnification. Sample counts were extrapolated to entire surface area of filter.

### Shotgun Metagenomes

Twenty liters of seawater were collected and kept cold for less than 4 hours during the commute back to the lab, where they were then immediately filtered. The sample was filtered sequentially through 3.0 µm and then 0.2 µm polycarbonate filters, using a peristaltic pump and 142 mm inline filter rigs. Filters were frozen at –80°C until further processing. DNA was extracted from the 0.2 µm and 3.0 µm filters with standard phenol chloroform methods. Extracted DNA was cleaned and purified with ethanol precipitation. Genomic DNA was processed with the Nextera XT Library Preparation kit. At least 3 replicate PCR amplification reactions were pooled with less than 10 cycles each to add Illumina adapter sequences and barcodes. Sequences were read on the Illumina NextSeq 500 platform with a NextSeq high output 300 cycle kit; 150 bases were read in each direction in addition to the 8 bp barcode used to differentiate reads from different size fractions. Metagenomic samples were sequenced at 207 and 156 million reads for the 0.2–3 µm, and >3 µm size fractions, respectively (Table S1).

### Hi-C Libraries

Five liters of water were collected and immediately treated with 1% formaldehyde. After 30 minutes, cross-linking reactions were quenched with the addition 10 mg/L of glycine. Within hours treated water was size-fractionated onto 3.0 µm and subsequent 0.2 µm polycarbonate filters. Material was then resuspended from the filters in 1x TE and pelleted by centrifugation. Pellets were frozen at –80°C and then delivered to Phase Genomics in Seattle, WA, for Hi-C library processing using established methods^39,72^. Briefly, restriction endonucleases with diverse AT content, Sau3AI and MluCI, were employed to digest formaldehyde fixed DNA in each sample. Digested ends were filled in, biotinylated, and blunt end ligated in a dilute environment to promote self-ligation. Biotinylated junctions were purified^39,52^ and sequenced on an Illumina HiSeq 2500, with 2 x 150 paired end reads, resulting in 80.6 million and 56.2 million reads for the 0.2–3.0 µm and the >3.0 µm libraries respectively (Table S1).

### Assembly and Hi-C independent Binning

Metagenomic reads were quality-filtered using Trimmomatic (v.0.33), targeting and extracting Illumina adapter sequences and reads with low phred scores^73^. Filtered reads were then assembled using MEGAHIT (v.1.1.1) on default settings, with multiple kmers^74^. The total assembled sequences were 1,147 Mb and 360 Mb with N50s of 1286 and 1454 for the 0.2–3 µm and the >3 µm libraries respectively (Table S1). Quality of assemblies was analyzed using QUAST^75^. Protein coding sequences were predicted using Prodigal^76^ (v2.6.3). For each library contig coverage was determined by recruiting reads to the assembly using Bowtie 2^77^ (v1.2.2). Assembled contigs in each size fraction were binned independently using MetaBAT 2^37^ (v2.15), with a minimum contig length of 1500 bp and default settings otherwise.

### Binning with Hi-C Reads

Hi-C read pairs recruited to two distinct contigs were used as proximity linkages and bins were generated with two separate algorithms that employed these linkages. First, ProxiMeta, processed by Phase Genomics in Seattle, WA, utilized adapted MetaPhase methods as previously described^39^. Secondly, the same Hi-C read pairs were run through an open-source pipeline, HiCBin^78^. In both methods contigs greater than 1000 bp were clustered together into bins using a normalized link frequency. In ProxiMeta, link frequency between two contigs is determined by mapping reads to an assembly using Bowtie2^77^ and then normalized between contigs based on the number of self-links for each contig, which considers contig size and frequency of enzyme digestion sites. With the HiCBin pipeline, HiC reads were mapped to assembled contigs using BWA-MEM with −5SP setting and HiCZin normalization was employed^79^. Because HiCBin requires coverage and taxonomy information for normalization, taxonomy assignments generated from a custom DIAMOND blastx (described below) were formatted with kaiju using the addTaxonNames script, with subsequent brief formatting modifications to meet HiCBin taxonomy requirements and include virus information^80^. Contig coverage information was determined using the MetaBAT 2 jgi_summarize_bam_contig_depths script. Normalized link frequencies were used by each method independently to cluster contigs into bins. ProxiMeta bins were further refined further to extract the largest connected component or giant component of its network graph (Figure S2) using CINNA and igraph libraries in R^81^.

The majority of contigs in each ProxiMeta giant component bin were assigned to the same HiCBin, but HiCBin typically lumped several ProxiMeta giant components together. Therefore, intersection bins were created by filtering ProxiMeta giant components to include only those contigs that went to the same HiCBin cluster and then combining those filtered giant components when the HiCBin method had combined them (Figure S3). Contigs not binned together by both methods were excluded from the final intersection bins. These intersection bins were then assessed for contamination and quality using CheckM. Bins which were >10% contaminated were split back into their ProxiMeta giant components, still filtered for contigs that all were binned in the same HiCBin cluster. Final bin names were IB_x.x_yyy, where IB means intersection bin, x.x is either 0.2 or 3.0 for size fraction library and yyy is the bin number.

### Bin Completeness and Contamination

Bins from every method were assessed for completion and quality using CheckM^82^ (v1.1.3) with the lineage_wf command. CheckM results were used to classify bins as high quality (HQ:>90% complete, <5% contaminated), medium quality (MQ:>50% complete, <10% contaminated), low quality (LQ:<50% complete, <10% contaminated) and contaminated (>10% contaminated) as previously described^83^. Two bins identified as *Picochlorum* were also assessed using BUSCO^52^ (v5.3.0) using the chlorophyta_odb10 lineage dataset (creation date: 2020-08-05, number of genomes: 16, number of BUSCO-derived single-copy orthologs: 1519).

### Taxonomic Identification of Assembled Contigs

A custom database was made by compiling three separate publicly available databases: MarDB, MMETSP, and Viral RefSeq. MarDB is a database of 8000 marine akaryote genomes^84^. The MarDB database was downloaded from the Marine Metagenomics Portal website in February of 2019^84^. The Viral RefSeq database was composed of all available viral genomes from RefSeq (release84). These databases were combined with the Marine Microbial Eukaryote Transcriptome Sequencing Project (MMETSP)^85^ with custom additions of zooplankton genomes. Each contig in the size-fractionated assemblies was identified using a translated blast with the fast local aligner DIAMOND^86^ (v0.9.24.125) using the Lowest Common Ancestor (LCA) parameter.

### Bin Taxonomy Assignment

Best LCA assignments and best phylum assignments were determined for each bin, by normalizing contig taxonomy assignments by length. Each bin was composed of contigs of varying sizes and best LCA taxonomic matches; therefore, the percentage of total DNA length of a bin identified as a specific taxon was used to determine the most likely taxonomy for every bin. Best LCA taxonomic identities were determined for each bin by the largest percentage of total DNA that was identified as that specific taxonomic ID. All LCA matches from the same Phylum were also summarized so that each bin had a best phylum, when possible, even if individual contigs could have been identified more specifically. If phylum assignment was not possible, highest order of taxonomic assignment possible was given.

### Identifying Viruses

Virus sequences were identified within each metagenome using several methods: VirSorter, VirFinder and DIAMOND blastx results. VirSorter^68^ (v1.0.5) was run within the CyVerse^69^ discovery environment and was applied to all contigs greater than 5 kb. Sequences returned as category 1 (most confident) or category 2 (very likely) were accepted as viral. VirFinder^87^ (v1.0.0) was run on Rstudio using default settings against all contigs greater than 5 kb. Contigs determined to be viral by either VirSorter, VirFinder or a virus best hit in the DIAMOND search against our custom database, as previously described, were considered potentially viral. All final intersection bins that contained more than 10 kb of potential viral sequence (27 bins) were investigated further using CheckV^88^ (v0.8.1); those identified as viral genomes (from 18 bins) were classified using DemoVir (https://github.com/feargalr/Demovir). Genes on these viral contigs were assessed for protein family hits on the pfam.xfam.org website^89^ and blasted against Viral RefSeq (release 211) using command line blastp^90^. Comparison of S-CAM7 virus genomes was generated in Easyfig^91^.

### Phylogenetic Analysis

Whole genome phylogenies for Planctomycetes, Chlorophyta and *Synechococcus* were constructed in PhyloPhlAn^92^ (v3.0.60) using settings –diversity medium and the config_file supermatrix_aa.cfg. Marker genes for each tree used the BUSCO-derived single-copy orthologs for each lineage (planctomycetes (571 genes), Chlorophyta (1519 genes) Synechococcales (788 genes)). For final phylogenetic trees the best protein model for the PhyloPhlAn generated concatenated alignment (LG+G4+F in all 3 cases) was found using ModelTest-NG^93^ (v0.1.7) and then used for a maximum likelihood search using RAxML-NG^94^ (v1.1) with 20 distinct starting trees. Trees were visualized and annotated using iTOL^95^ (v6.7). Average nucleotide identity (ANI) values were calculated using FastANI^96^ (v1.33).

### Data Deposition

Sequences have been deposited in GenBank under BioProject PRJNA992579. Hi-C reads SRR25203940-SRR2520394, short reads SRR25203943-SRR25203944.

## Acknowledgements

This work was supported by NSF DEB-1542240 to GR. CR was also supported by a National Science Foundation Graduate Research Fellowship. We thank I. Liachko and Phase Genomics for processing and sequencing of Hi-C libraries. Additionally we would like to thank the ORCA team and crew of the R/V Mackinaw for assistance in collecting samples and sharing chemical data as well as J. Deming for helpful comments on the manuscript.

CR and GR conceived the study. CR, CM and GR performed analyses. CR and GR wrote the paper.

The authors declare no competing interests.

